# In Vivo Tissue Distribution of Microplastics and Systemic Metabolomic Alterations After Gastrointestinal Exposure

**DOI:** 10.1101/2023.06.02.542598

**Authors:** Marcus M. Garcia, Aaron S. Romero, Seth D. Merkley, Jewel L. Meyer-Hagen, Charles Forbes, Eliane El Hayek, David P. Sciezka, Rachel Templeton, Jorge Gonzalez-Estrella, Yan Jin, Haiwei Gu, Angelica Benavidez, Russell P. Hunter, Selita Lucas, Guy Herbert, Kyle Joohyung Kim, Julia Yue Cui, Rama Gullapalli, Julie G. In, Matthew J. Campen, Eliseo F. Castillo

## Abstract

Global plastic use has consistently increased over the past century with several different types of plastics now being produced. Much of these plastics end up in oceans or landfills leading to a substantial accumulation of plastics in the environment. Plastic debris slowly degrades into microplastics (MPs) that can ultimately be inhaled or ingested by both animals and humans. A growing body of evidence indicates that MPs can cross the gut barrier and enter into the lymphatic and systemic circulation leading to accumulation in tissues such as the lungs, liver, kidney, and brain. The impacts of mixed MPs exposure on tissue function through metabolism remains largely unexplored. To investigate the impact of ingested MPs on target metabolomic pathways, mice were subjected to either polystyrene microspheres or a mixed plastics (5 µm) exposure consisting of polystyrene, polyethylene and the biodegradability and biocompatible plastic, poly-(lactic-co-glycolic acid). Exposures were performed twice a week for four weeks at a dose of either 0, 2, or 4 mg/week via oral gastric gavage. Our findings demonstrate that, in mice, ingested MPs can pass through the gut barrier, be translocated through the systemic circulation, and accumulate in distant tissues including the brain, liver, and kidney. Additionally, we report on the metabolomic changes that occur in the colon, liver and brain which show differential responses that are dependent on dose and type of MPs exposure. Lastly, our study provides proof of concept for identifying metabolomic alterations associated with MPs exposure and adds insight into the potential health risks that mixed MPs contamination may pose to humans.

## INTRODUCTION

Over the past 50 years, global plastic production has grown exponentially. To date, approximately 350 million metric tons of plastic are produced globally every year (Roland Geyer et al. 2017). Much of this plastic ends up in landfills or oceans where it may take several hundred years to degrade depending on composition and environmental factors (Bajt 2021). Exposure to light, heat, moisture and microbes degrades plastic debris into microplastics (MPs), defined as plastic particles smaller than 5mm (Anderson et al. 2017). MPs have become ubiquitous throughout our environment and exposure to humans and animals occurs via ingestion or inhalation (Cheng et al. 2022; El Hayek et al. 2023; R. Geyer et al. 2017; Schymanski et al. 2018). Multiple studies have reported MP detection in salt water, fresh water, farming soils, and crops used for both animal and human consumption (Deng et al. 2017; Nizzetto et al. 2016), (Hirt and Body-Malapel 2020; Priya et al. 2022; Ren et al. 2021; Udovicki et al. 2022; Vivekanand et al. 2021). A 2019 review of over 50 existing studies on MPs suggests that consumption of common foods and beverages results in humans ingesting approximately 5 grams of plastic per week (N Biguad 2019). It is currently estimated that by 2050, approximately 12 billion metric tons of plastic wastes will be released into the environment by bioturbation, atmospheric deposition, sewage irrigation, and landfills (R. Geyer et al. 2017; Nizzetto et al. 2016; Ren et al. 2021; Rillig et al. 2017). With an estimated 3.2 (and growing) metric tons of MPs being released into the environment via commercial and household activities every year, MPs exposure is now unavoidable (Julien Boucer 2017).

In 2019 the World Health Organization (WHO) released a statement that, based on the limited available evidence, exposure to MPs poses a low concern for human health (Organization 2019). However, subsequent reports of cellular and biochemical toxicity of MPs have made it clear that deleterious interactions between MPs and biological systems are dose-dependent, meaning that the impacts of MPs exposures will increase with time (Hayek et al. 2022). Ingestion is believed to be the most common route of MPs exposure. Thus, it is no surprise MPs induce gut microbiota dysbiosis, in both mice and zebrafish (Jin et al. 2018; Jin et al. 2019; Lu et al. 2018; Zhao et al. 2021). Gut dysbiosis is linked to numerous inflammatory and metabolic diseases (Aron-Wisnewsky et al. 2019; Boursier et al. 2016; Joossens et al. 2011; Kaur et al. 2011; Sata et al. 2020; Tamboli et al. 2004) and studies in zebrafish exposed to MPs have been shown to induce intestinal injury and inflammation (Jin et al. 2018; Limonta et al. 2019; Y. Lu et al. 2016; Qiao et al. 2019a; Qiao et al. 2019b; Wan et al. 2019). In contrast to these reports, other studies have concluded MPs caused no intestinal histological damage in the colon of mice (Stock et al. 2019; Sun et al. 2021). In humans, recent evidence show MPs are abundant in colons collected after a colectomy (Ibrahim et al. 2021) and increased in the stool of individuals with inflammatory bowel disease (Yan et al. 2022); however, the effects of MPs on GI health is still being deciphered. Additionally, studies in marine animals, rodents, and human cell cultures have shown that MPs can alter cellular bioenergetics, induce oxidative stress and inflammation in various immune cells and extraintestinal organs (Cheng et al. 2022; Merkley et al. 2022; Qiao et al. 2019b).

Exposing animals to MPs via the oral gastric route leads to the dissemination of MPs outside of the intestine, specifically the liver (Deng et al. 2017). Additionally, several studies have shown MPs ingestion results in their accumulation in tissues such as the liver and kidney (Cheng et al. 2022; Hirt and Body-Malapel 2020; Yifeng Lu et al. 2016; Luo et al. 2019; Vethaak and Legler 2021). This distribution into the liver caused metabolic alterations (Cheung et al. 2018; Jin et al. 2019; Jing et al. 2022). These studies highlight how MPs can cross the intestinal barrier and can cause extraintestinal manifestations. However, there is a lack of evidence regarding the effects of a combination of microplastics. The studies conducted so far have solely focused on investigating the impacts of polystyrene microplastics, nor have they examined the metabolic changes that can take place in organs that have a direct interaction with the gut, such as the kidney and brain. As we begin to understand the environmentally-relevant sizes, types, and doses of MPs humans are ingesting (Blackburn and Green 2022; Kieran D. Cox et al. 2019; Schwabl et al. 2019; Wright and Kelly 2017), researchers are able to further assess the effects of MPs using environmentally-relevant doses, sizes, and types. To date, the systemic impacts of MP ingestion on metabolomic pathways in systemic organs have not been studied utilizing environmentally relevant doses and mixtures. Therefore, we set out to understand how polystyrene MPs (5 µm) and mixed MPs (5 µm) exposure consisting of polystyrene, polyethylene, and poly-(lactic-co-glycolic acid) at environmentally-relevant doses cross the intestinal barrier and alter metabolism in the colon, liver and brain. Specifically, we evaluated the systemic distribution and metabolomic impacts of polystyrene MPs and mixed plastics ingestion in mice after oral-gastric exposure. While not reflective of the myriad environmental MPs, plastic microspheres are a research model that controls for size, shape, and composition, while eliminating the contributions of endotoxin, pyrogens, metals, or other contaminants that may have interactive and confounding effects. Microscopic visualization and Raman spectroscopy were used to evaluate MPs accumulation in the colon and translocation to the liver and brain, assess composition post-exposure, and identify any physical or chemical changes associated with biological degradation. Targeted and untargeted metabolomic profiling was then used to identify functional responses within the colon, liver, and brain. These studies confirm systemic translocation of plastics and metabolomic disruption in a mammalian model. This study is also one of the first to evaluate impacts of not only polystyrene MPs but also a mixed plastics model at an equivalent dose to what humans are estimated to consume per week.

## METHODOLOGY

### Animal Model

Male and female C57BL/6 mice (8-12 weeks of age at the beginning of the study) were obtained from Taconic Biosciences (Rensselaer, New York). Animals were housed in an Association for Assessment and Accreditation of Laboratory Animal Care (AAALAC)-approved facility at the University of New Mexico Health Sciences Center. Animals were maintained at constant temperature (20–24°C), relative humidity (30%–60%), and on a 12-h light/dark cycle throughout the study. Animals were provided with normal chow and water *ad libitum*. All experiments were approved by the Institutional Animal Care and Use Committee of the University of New Mexico Health Sciences Center, in accordance with the National Institutes of Health guidelines for use of live animals. The University of New Mexico Health Sciences Center is accredited by the American Association for Accreditation of Laboratory Animal Care.

### Exposure, Tissue Digestion, and Microplastic Isolation

Mice were exposed twice a week to MP via oral gastric gavage over a four-week period at 0 mg/week (n=8), 2 mg/week (n=8), and 4 mg/week (n=8) of 5 µm microspheres from Degradex® Phosphorex (Hopkinton, MA). Two different microsphere exposures were used for each dose group, for a total of six groups. The two different microsphere exposures were 1) polystyrene (PS) microspheres and 2) a mixed plastics treatment consisting of polystyrene (PS), polyethylene (PE), and poly (lactic-co-glycolic acid) (PLGA) microspheres. Prior to gavage, all microspheres were stored based on the manufacturer’s recommendations in a 10 mg/ml concentration at 4°C until exposure. After four weeks, the mice were euthanized and exsanguinated, then systemically perfused with ice cold saline to ensure removal of blood from major organs. Serum, brain, liver, kidney, and colon were isolated and stored in glass vials to prevent MP contamination from the postmortem procedures.

Digestion of the prefrontal cortex of the brain, left lobe of liver, and sagittal cross section of kidney tissues was performed using 3 x the sample volume of 10% potassium hydroxide (KOH) prepared using deionized water. Samples were incubated at 40°C with agitation for 72 hours. Samples were ultra-centrifuged (Thermo Scientific Sorvall WX+) at 30,000 x g for 4 hours to isolate MPs into a pellet. The supernatant was removed and the pellet was washed with 100% ethanol (EtOH) and centrifuged at 15,000 x g for 10 minutes, followed by removal of excess EtOH. The process was repeated three times. After washing, the samples were re-suspended in EtOH and stored at 4°C in glass tubes for processing.

To prevent adulteration from airborne and procedural contaminants, we used the following preventative measures: (i) Latex gloves were worn during all experiments along with 100% cotton lab coats, (ii) all reagents were prepared using ultrapure water, (iii) all samples were stored in glass vials (DWK Life Sciences), (iv) all surfaces were covered in absorbent bench coat and cleaned with 70% ETOH prior to experiments, (v) use of plastic containing equipment was kept to a bare minimum, (vi) and all tools were cleansed and autoclaved prior to use.

### Visualization and Spectroscopic Characterization of Microplastics

Light Microscopy: Samples of isolated plastics were stored in 100% EtOH in glass tubes for a minimum of 24 hours at 4°C before imaging. Slides were prepared by adding 50 μl of sample and MPs were identified and imaged via polarized light microscopy using an Olympus BX51 microscope. An ultraviolet light source was used to verify that the remaining solids were plastic in nature based on autofluorescence.

Raman Spectroscopy: Raman Spectroscopy analysis was processed on a WITec Alpha 300R Confocal Raman microscope with a 532nm laser on isolated MPs from brain to confirm that MPs of interest were polystyrene. Additionally, pristine 5µm polymeric microspheres that underwent 10% Potassium hydroxide (KOH) digestion protocol were analyzed to determine if MPs alteration occurs based on biological or chemical degradation. A series of peaks specific to the materials were generated. Substance-specific peaks from an Infrared and Raman Characteristic Group Frequencies library were compared to the generated data to identify the materials.

X-ray Photoelectron Spectroscopy (XPS) Analysis: A Kratos Ultra DLD spectrometer with a monochromatic Al Kα source operating at 150W (1486.6 eV) was used to perform XPS measurements. The spectrometer was operating at a pressure of 5 x 10^-9^ Torr. Low energy electrons were used to accomplish a charge compensation, and all spectra charges were referenced and adjusted by the C1s region to 285 eV. All high-resolution C1s and survey spectra were acquired at pass energies of 160 and 20 eV and processed using CasaXPS software.

### Metabolomic Analysis of Colon, Liver, and Brain

Reagents used in this study were all liquid chromatography–mass spectrometry (LC-MS) grade and all standard components for measuring metabolites were purchased from both Sigma-Aldrich (Saint Louis, MO) and Fisher Scientific (Pittsburg, PA). Acetonitrile, ammonium acetate, acetic acid, and methanol (MeOH) were purchased from Fisher Scientific. Ammonium hydroxide was purchased from Sigma Aldrich. This study used deionized water that was produced by an in-house water purification system from EMD Millipore (Billerica, MA). Phosphate Buffered Saline (PBS) was purchased through GE Healthcare Life Sciences (Logan, UT).

### Tissue preparation

To prepare each tissue sample for analysis, approximately 20 mg of the tissue of interest was homogenized in an Eppendorf tube using a Bullet Blender homogenizer (Next Advance, Averill Park, NY). Each sample was homogenized in 200 µL MeOH: PBS (a 4:1 volume to volume dilution containing 1,810.5 μM ^13^C_3_-lactate and 142 μM ^13^C_5_-glutamic Acid). After completing the initial homogenization, an additional of 800 µL MeOH:PBS was added and samples were vortexed for 10 seconds. Thereafter, samples were stored at -20°C for 30 minutes, transferred to an ice bath and sonicated for 30 minutes. Centrifugation at 14,000 rpm was then performed at 4°C for 10 minutes and 800 µL supernatant was transferred to a new Eppendorf tube. Drying of all samples was performed using a CentriVap Concentrator (Labconco, Fort Scott, KS) under vacuum and all residue obtained was reconstituted in 150 μL 40% PBS and 60% acetonitrile prior to MS analysis. A portion of All study samples were pooled together to obtain for quality control (QC) samples.

### Targeted Metabolomics

The Agilent 1290 UPLC-6495 QQQ-MS system was utilized to perform LC-MS/MS experiments targeting metabolites in the tryptophan metabolic pathway (Scieszka et al. 2022). There were 28 metabolites targeted in this evaluation, including 3-hydroxy anthranilic acid, 3-hydroxykynurenine, 3-Indolepropionic acid, 5-Hydroxytryptophan, ADP ribose, Anthranilic acid, HIAA, Indole, Indole-3-acetic acid, Indole-3-lactic acid, Indole-3-pyruvic acid, Kynurenic acid, L-kynurenine, Melatonin, NAD, NADH, N’-Formylkynurenine, Nicotinamide, nicotinamide mononucleotide, nicotinamide riboside, Nicotinic acid, Nicotinic acid adenine dinucleotide, Nicotinic acid mononucleotide, Quinolinic acid, Serotonin, Tryptamine, Tryptophan, and Xanthurenic acid. Each sample was injected using a volume of 4 µL and analyzed in positive ionization mode. Chromatographic separations were conducted through a Waters XBridge BEH Amide column in hydrophilic interaction chromatography (HILIC) mode. The flow rate was set to 0.3 mL/min, and the auto-sampler temperature and column compartment were maintained at 4°C and 40°C, respectively. The mobile phase consisted of two solvents: the first solvent containing 10 mM ammonium acetate and 10 mM ammonium hydroxide in 95% H_2_O and 5% acetonitrile, and the second solvent containing 10 mM ammonium acetate and 10 mM ammonium hydroxide in 95% acetonitrile and 5% H_2_O. An isocratic elution of 90% of the second solvent was performed for 1 minute, followed by a decrease to 40% at timepoint t=11 minutes for 4 minutes, then gradually increasing back to 90% at timepoint t=15 minutes. The mass spectrometer was equipped with an electrospray ionization (ESI) source to acquire targeted data in multiple-reaction monitoring (MRM) mode. The LC-MS system employed the Agilent MassHunter workstation software (Santa Clara, CA), while the Agilent MassHunter Quantitative Data Analysis Software was used to integrate all extracted MRM peaks during analysis. *Untargeted LC-MS Metabolomics*

The untargeted LC-MS metabolomics analysis was conducted using a Thermo Scientific Vanquish UHPLC system with a Thermo Scientific Orbitrap Exploris 240 MS (Waltham, MA). Duplicate samples of 10 µL were analyzed in negative ionization mode, and an additional 4 µL was injected for positive ionization mode. A Waters XBridge BEH Amide column (150 x 2.1 mm, 2.5 µm particle size, Waters Corporation, Milford, MA) was used for chromatographic separation in hydrophilic interaction liquid chromatography (HILIC) mode, with a flow rate of 0.3 mL/min and auto-sampler temperature was kept at 4°C with the column compartment set to 40°C. The LC conditions were the same as those in targeted metabolomics described above. The mass spectrometer collected untargeted data at 70 to 1,000 m/z using an electrospray ionization (ESI) source. MS spectra peaks were identified using approximately 600 aqueous metabolites of in-house chemical standards, and compared several MS databases, including the HMDB library, Lipidmap database, METLIN database, and commercial databases. Limits of 5 ppm for mass accuracy and 1,000 for absolute intensity threshold were set for MS data extraction. Data annotation was based on isotopic pattern, retention time, exact mass, and MS/MS fragmentation patterns. Thermo Scientific Compound Discoverer 3.3 software was used for data processing of aqueous metabolomics data, and peak picking, alignment, and normalization was used for untargeted data. Quality control (QC) pools were established based on the coefficient of variation (CV) < 20% and signals showing up in > 80% of all samples to ensure high-quality data for analysis.

### Statistical Analysis

All samples were analyzed and compared against untreated control mice using a 10% False Discovery Rate, where appropriate. Pathway analysis and volcano plots were developed using MetaboAnalyst software 5.0 (www.metaboanalyst.ca). Samples were normalized by sum against QC pools. Log transformation was performed and all data was mean centered. P-value threshold was set to 0.05 with equal variance and 2.0-fold change threshold.

## RESULTS

### Visualization of Systemic Microplastic Translocation

To determine whether orally dosed microspheres could be translocated from the digestive system, polarized light microscopy was performed on serum, brain, liver, and kidney samples isolated from mice exposed to 0, 2, 4 mg/week polystyrene and mixed plastic (polystyrene, poly(lactic-co-glycolic acid), and polyethylene) microspheres for four weeks. The presence of MPs were observed in the serum and in all three isolated tissues (**Figure 1-2**). Although not fully quantifiable with this visualization method, MPs were readily more apparent in liver samples (**Figure 1C**) compared with brain and serum, with far fewer MPs observed in the kidneys. These observations confirm that ingested MPs are able to translocate across the gut epithelium into the systemic circulation and accumulate differentially in the assessed organs.

**Figure 1.**
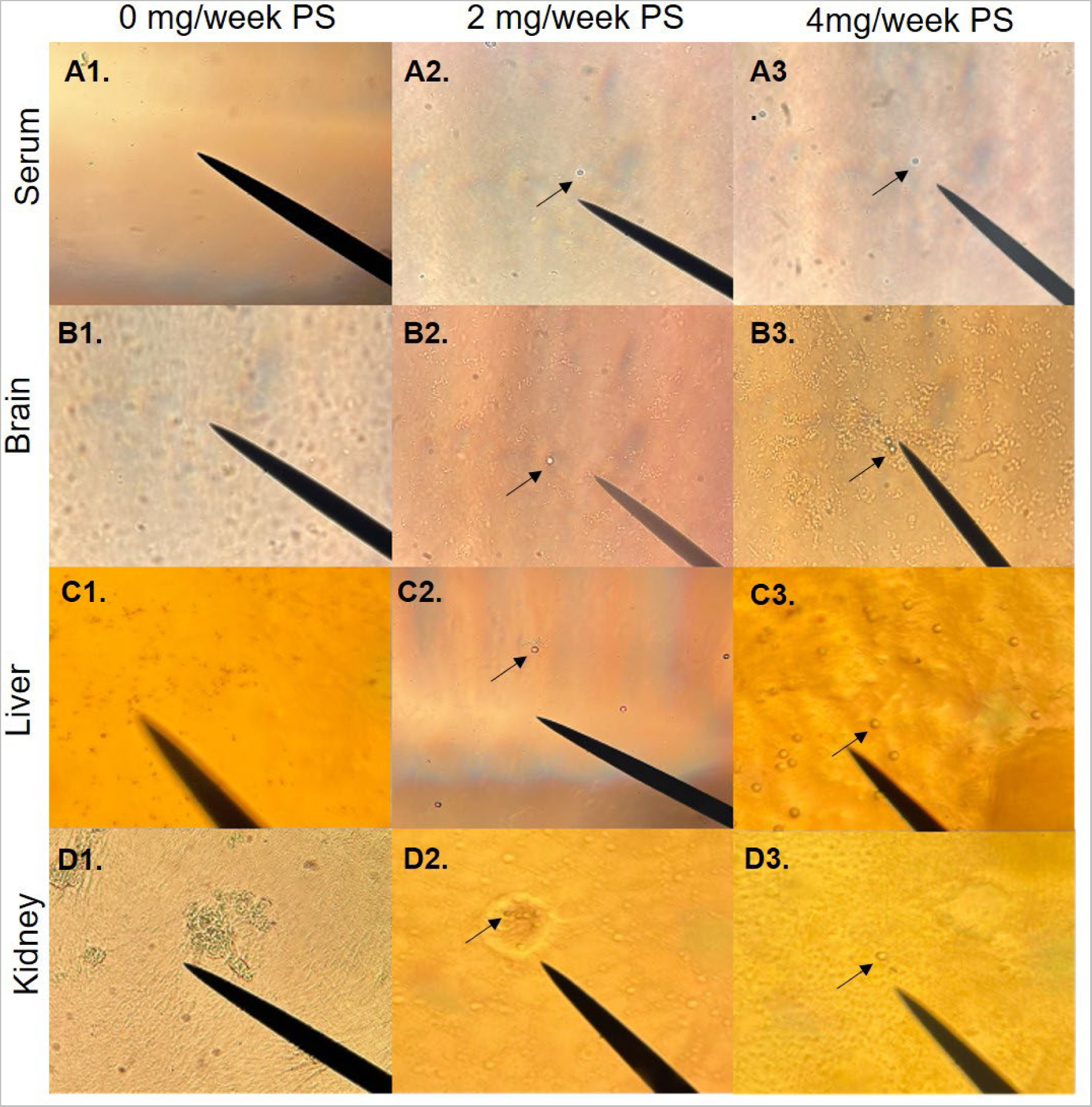
Oral exposure to microplastics for 4 weeks causes accumulation in the serum, brain, liver, and kidney of mice. The black arrow indicates A1-A3. 5 µm polystyrene microspheres in serum (20x), B1-B3. 5 µm polystyrene microspheres in brain (20x), C1-C3. 5 µm polystyrene microspheres in liver (40x), and D1-D3. 5 µm polystyrene microspheres in kidney (40x). Mice were exposed twice a week for 4 weeks to 2 or 4 mg 5 µm polystyrene microspheres via oral gavage. Images are representative of n=8.

**Figure 2.**
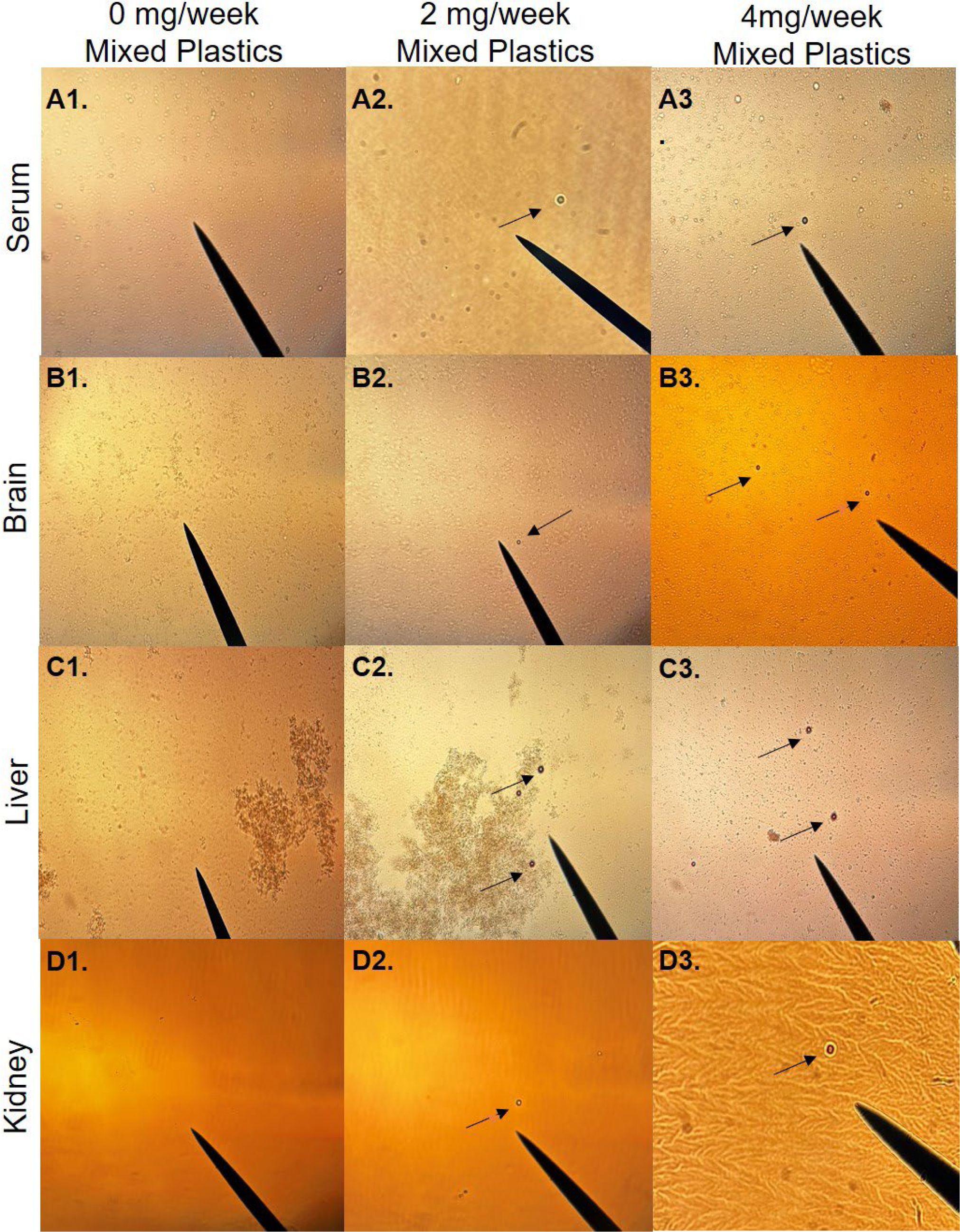
Oral exposure to microplastics for 4 weeks causes accumulation in the serum, brain, liver, and kidney of mice. The black arrow indicates E1-E3. 5 µm Mixed MPs (PS, PE, PLGA) microspheres in serum (20x), F1-F3. 5 µm Mixed MPs (PS, PE, PLGA) in brain (20x), G1-G3. 5 µm Mixed MPs (PS, PE, PLGA) in liver (20x), and H1-H2. 5 µm Mixed MPs (PS, PE, PLGA) in kidney (40x), H3 5 µm Mixed MPs (PS, PE, PLGA) in Kidney (40x). Mice were exposed twice a week for 4 weeks to 2 or 4 mg 5 µm Mixed MPs (PS, PE, PLGA) microspheres via oral gavage. Images are representative of n=8.

Using Raman Spectroscopy and XPS analysis, we wanted to validate the brain MPs isolated and viewed under polarized light microscopy were truly polystyrene microspheres. The Raman spectra of the original 5 µm polystyrene microspheres was consistent with those particulates found in brain isolates (**Suppl. Figure 1A**), and further matched a library polystyrene standard spectra, validating that the systemically translocated and recovered microspheres were polystyrene (**Figure 1B2, B3**). The microsphere sample recovered from brain isolate, however, did show peak shift alterations as compared to fresh polystyrene, potentially indicating modification of the surface chemistry or accumulation of other biochemicals (*i.e.,* a corona effect). This observation led us to question whether the alteration was due to the KOH digestion or biological degradation. To address this, we compared the recovered polystyrene microspheres to naïve polystyrene microspheres digested with KOH under the same conditions used in our isolation protocol (**Suppl. Figure 1B**). The Raman spectra for the naïve polystyrene microspheres subjected to our isolation protocol revealed that the KOH digestion did not alter the structure of the polystyrene microspheres. This finding indicates that polystyrene microspheres may be altered biologically during translocation.

**Figure 3.**
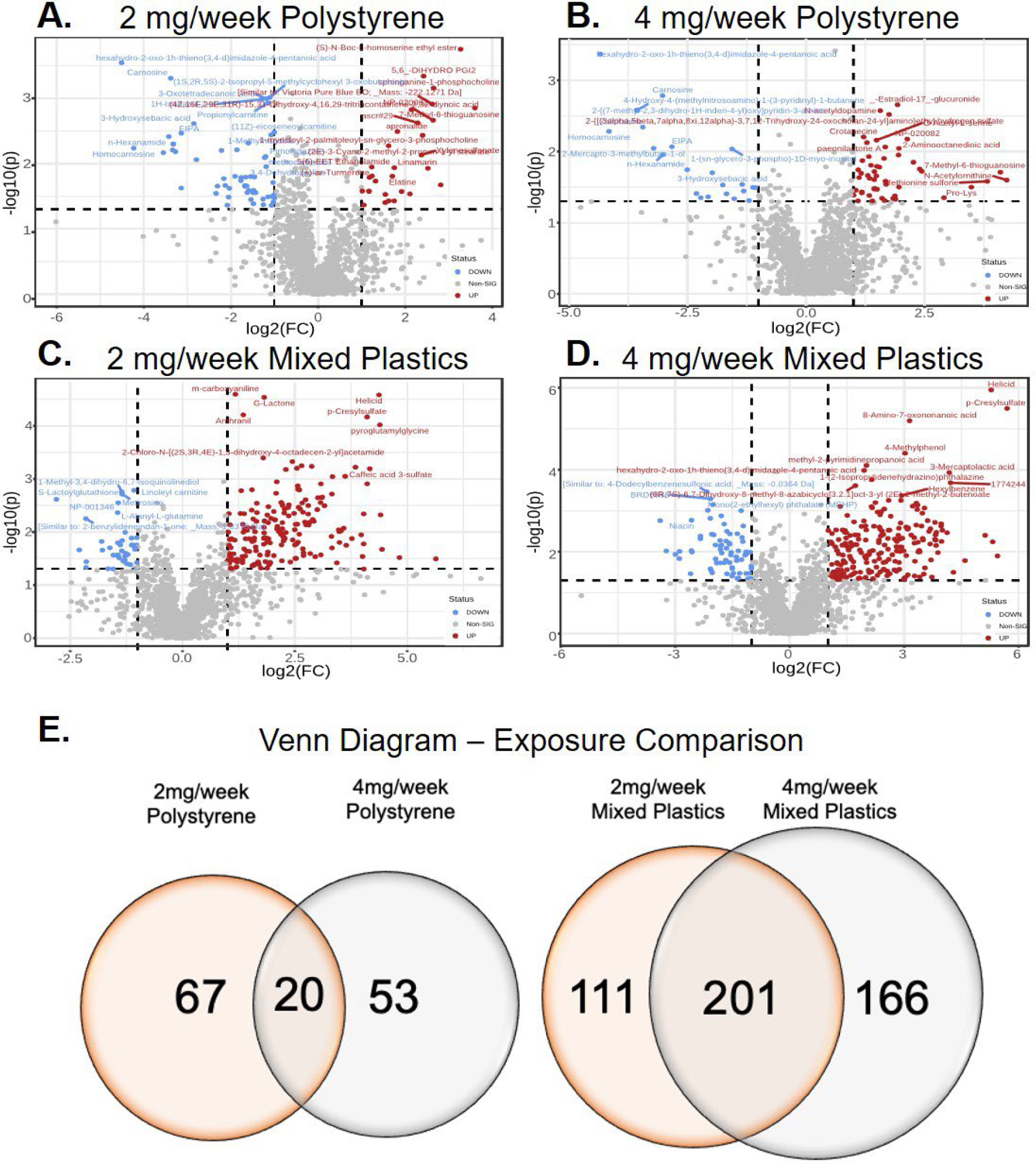
Untargeted metabolomic analysis in colon tissue of mice exposed to A) 2 mg/week polystyrene B) 4 mg/week polystyrene, C) 2 mg/week mixed plastics, or D) 4 mg/week mixed plastics. Data plotted as log(2) fold change (p = 0.05). E) Venn diagram representing the significantly changed metabolites following microplastic exposures (p < 0.05 as compared to control). Exposures (5 µm microspheres): 2 x week for 4 weeks oral gavage; Mixed plastics: polystyrene, polyethylene, and poly-(lactic-co-glycolic acid); n=8 per group.

Given the liver had the highest concentrations of MPs we wanted to investigate the chemical and surface composition of polystyrene microspheres recovered from exposed mice compared to naïve polystyrene microspheres using XPS survey scan mode (**Suppl. Figure 1C-E**). XPS analysis revealed increased surface potassium and nitrogen in the recovered microspheres, consistent with a biochemical adherence or interaction. XPS analysis also identified fluorine on the surface of the microspheres isolated from liver tissue. It was subsequently determined that this fluorine peak resulted from the surface adsorption of isoflurane, which was used as a general anesthetic prior to euthanasia in our studies. This fluorine peak may serve as a useful indicator that plastics were obtained from an anesthetized subject, as opposed to derived from *ex vivo* processing or storage of tissue samples in polystyrene containers.

### Untargeted Metabolomic and Pathway Analysis in the Colon

A few studies have performed both targeted and untargeted metabolomics in the serum (Deng et al. 2017; Jin et al. 2019; Zhao et al. 2023), liver (Wang et al. 2022), and stool (Schwabl et al. 2019) of micro- and nanoplastic-exposed mice; however, these mice were exposed to a single-type of MP (or NP). Given humans are exposed to a plethora of plastics, we set out to identify, quantify, and compare the colon metabolome of mice exposed to both doses of PS or mixed plastics after a 4-week exposure. Untargeted metabolomics was performed on colonic tissue metabolites in response to environmentally relevant oral MP exposure. Volcano plots for each exposure group showed both significantly increased and decreased metabolites when compared to control mice (**Figure 3A-D**). Following the 4-week exposures, 140 metabolites in the polystyrene exposed group and 478 metabolites in the mixed plastics exposed group were significantly altered (p < 0.05; **Figure 3E**). We observed that 67 metabolites were uniquely altered in the 2 mg/week polystyrene group and 53 metabolites were uniquely altered in the 4 mg/week polystyrene group, with 20 (14.3%) significantly altered metabolites occurring in both dose groups (**Figure 3E**). The mixed plastics exposure groups showed a much higher metabolomic alteration response, with 111 uniquely altered metabolites in the 2 mg/week exposure group and 166 uniquely altered metabolites in the 4 mg/week group. The mixed plastic groups shared 201 (42.0%) altered metabolites (**Figure 3E**).

To further understand these changes in metabolites, metabolomic pathway analysis were performed. Pathway analysis of metabolomic alterations in colons from mice exposed to 2 mg/week and 4 mg/week polystyrene compared to controls revealed significant alterations (p<0.05) in shared pathways contributing to i) biotin metabolism, ii) histidine metabolism, iii) β-alanine metabolism, iv) arginine and proline metabolism, v) cytochrome P450 xenobiotic metabolism, and vi) porphyrin and chlorophyll metabolism (Figure 4A, B). Interestingly, both mixed plastics exposed groups exhibited some overlap in metabolomic pathway when compared to the polystyrene only groups including i) porphyrin and chlorophyll metabolism, ii) primary bile acid biosynthesis, iii) arginine and proline metabolism, iv) arachidonic acid metabolism, and v) the pentose phosphate pathway (Figure 4A-D). Whereas the 2 mg/week and 4 mg/week mixed plastics groups only shared changes in pathways linked to the i) primary bile acid biosynthesis and ii) the biotin synthesis pathway. These data demonstrate a single MP exposure versus a mixed plastic exposure have varying effects in the colon.

**Figure 4.**
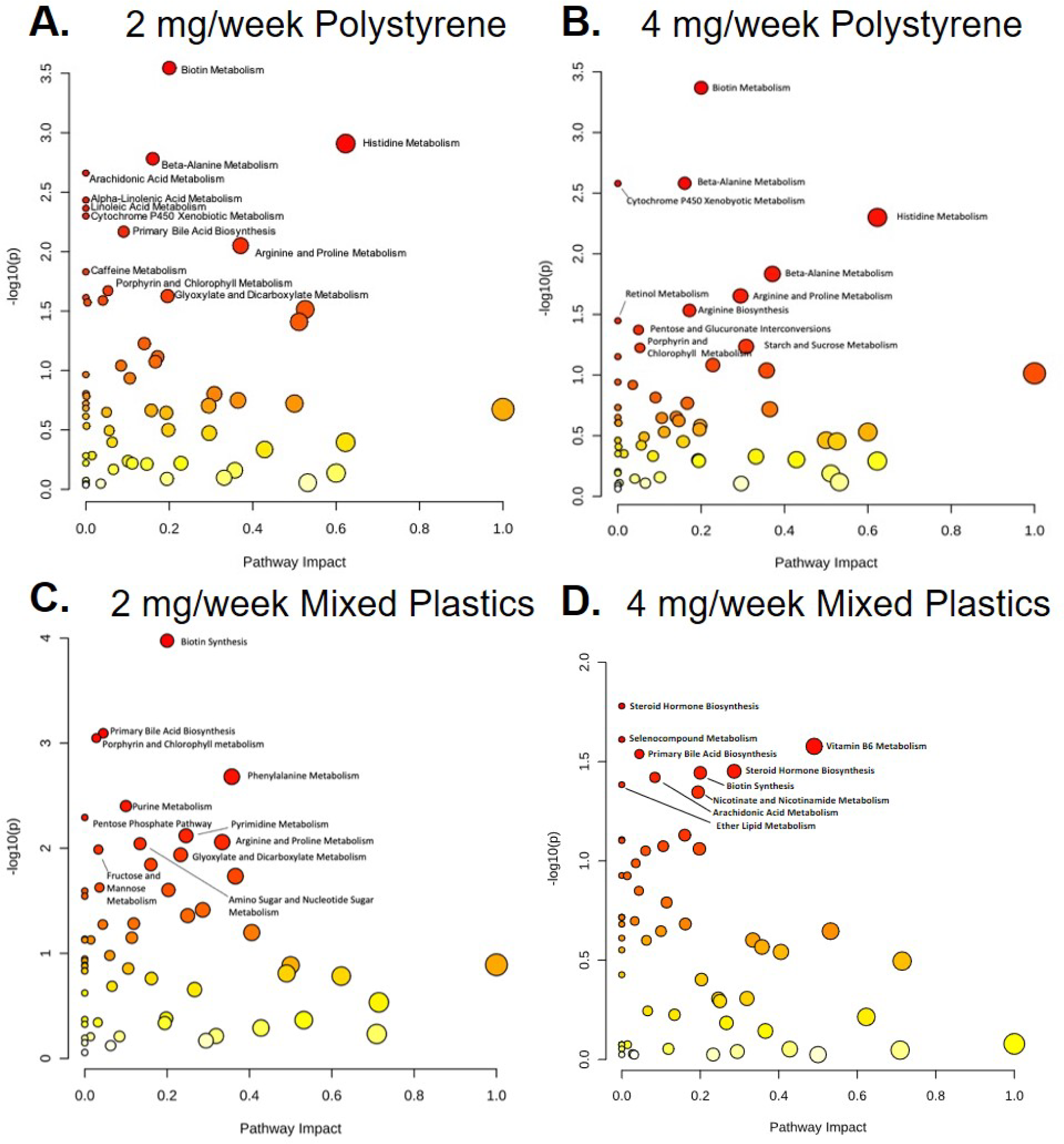
Metabolomic Pathway Analysis of alterations in the colon following oral MP exposure in mice exposed to A) 2 mg/week polystyrene, B) 4 mg/week polystyrene, C) 2 mg/week Mixed Plastics, or D) 4 mg/week Mixed Plastics. Exposures (5 µm microspheres): 2 x week for 4 weeks oral gavage; Mixed plastics: polystyrene, polyethylene, and poly-(lactic-co-glycolic acid); p<0.05 as compared to control; n=8 per group.

### Nicotinate and Nicotinamide Metabolic Pathway in the colon

High oxygen consumption (an oxidative metabolic state) in the colon is critical for intestinal homeostasis (Furuta et al. 2001; Litvak et al. 2018). Fatty acid oxidation and oxygen consumption through oxidative phosphorylation help sustain intestinal epithelial hypoxia. This prevents oxygen from crossing the barrier and subsequently preserves an obligate anaerobic microbial community as well as prevent gut microbial dysbiosis (Byndloss et al. 2017; Litvak et al. 2017; Litvak et al. 2018; Rivera-Chavez et al. 2017). Nicotinamide adenine dinucleotide (NAD) can be both an electron and hydrogen carrier in the process of oxidative metabolism during fatty acid oxidation and subsequent tricarboxylic acid (TCA) cycle to produce ATP (Xue et al. 2022). NAD and its metabolites can also function as critical regulators for cells to adapt to environmental changes in various tissues across the body (Xie et al. 2020). Therefore, we performed targeted metabolomic analysis of 28 metabolites associated with nicotinate and nicotinamide metabolic pathway (Suppl. Figure 2A-D). Analysis of colon tissue isolated from mice exposed to polystyrene only revealed upregulation of 5-hydroxytryptophan for both doses and downregulation of melatonin in the 2 mg/week dose group (Suppl. Figure 2A, B). The mixed plastics group shared an upregulation of kynurenic acid, xanthurenic acid, melatonin, and N’-formylkynurenine metabolites (Suppl. Figure 2C, D). A significant downregulation of the nicotinamide and ADP ribose metabolites was noted in the 2 mg/week group mixed plastic group while the 4 mg/week mixed plastics group exhibited a significant downregulation of NAD, indole, and nicotinic acid, and nicotinic acid adenine dinucleotide. In summary, we established that our findings indicate mixed plastics have a distinct effect on the colon compared to single-type plastics as revealed by our data.

### Oral gastric exposure of MP alters the liver metabolome

Numerous papers have shown gut exposure to a single-type of MP results in the accumulation of particles in the liver likely due to the gut-liver axis (Shi et al. 2022; Wang et al. 2022). Given these studies also examined a single plastic (Chen et al. 2022; Cheng et al. 2022; Jiang et al. 2023; Shi et al. 2022; Tu et al. 2023; Wang et al. 2022), we were particularly interested in metabolomics changes from mice exposed to the single PS plastics compared to the mixed plastic exposure. Therefore, untargeted metabolomics was performed on liver tissues from all microplastic exposed animals compared to non-exposed mice. Volcano plots for each exposure and dose group showed patterns of metabolite changes within each group (Figure 5). We observed 188 metabolites in the polystyrene exposed group and 137 metabolites in the mixed plastics exposed group were significantly changed due to MP exposure (p < 0.05). We observed that 129 (68.6%) metabolites were uniquely altered in the 2 mg/week polystyrene group and 16 (8.5%) metabolites were uniquely altered in the 4 mg/week polystyrene group, with 43 (22.9%) significantly altered metabolites in common between the two different dose of PS groups (Figure 5E). In contrast, the mixed plastics exposure groups showed a much higher metabolomic alteration response, with 20 (14.6%) uniquely altered metabolites in the 2 mg/week exposure group and 44 (32.1%) uniquely altered metabolites in the 4 mg/week group, and 73 (53.3%) altered metabolites shared amongst both mixed plastic doses.

**Figure 5:**
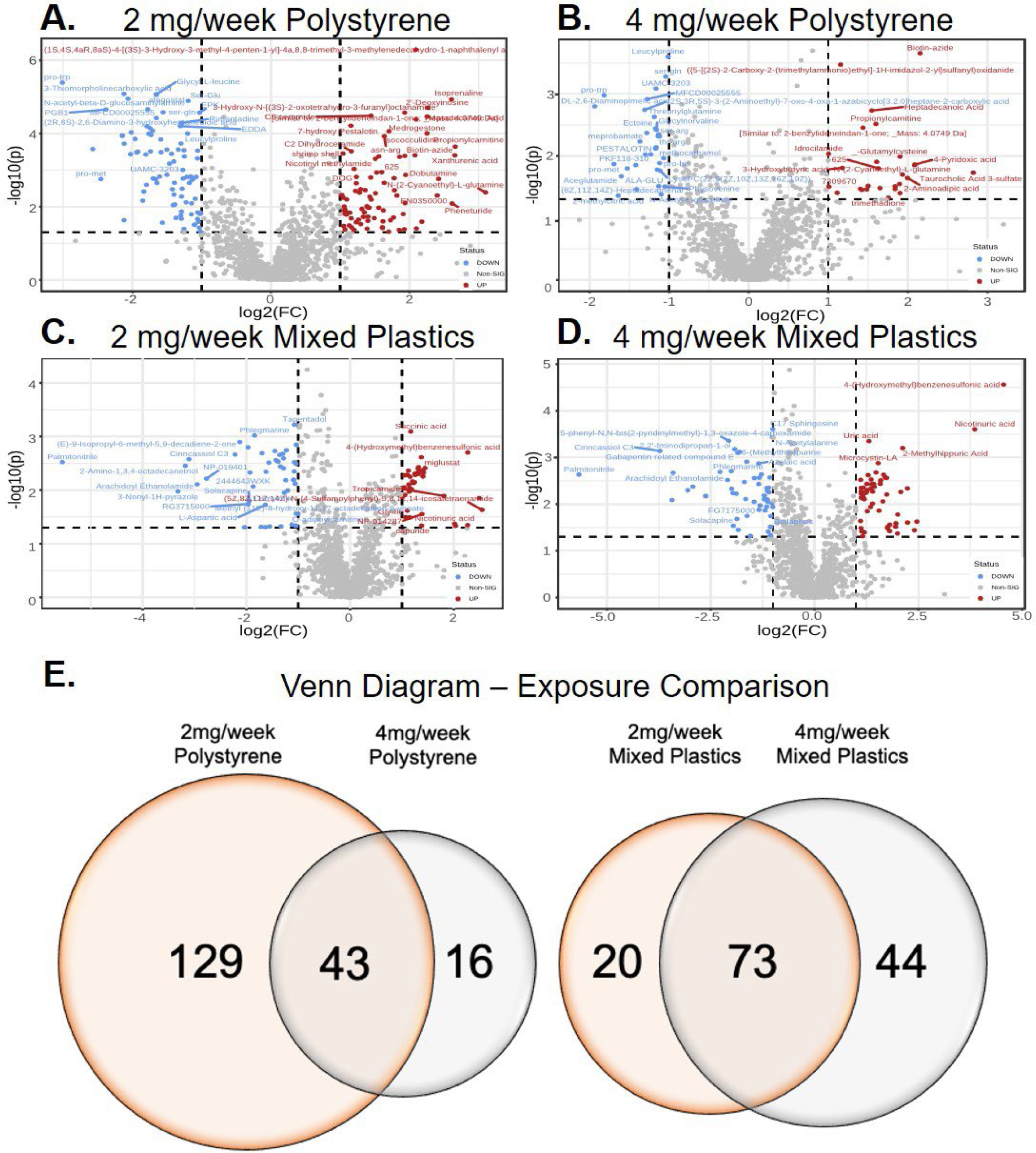
Untargeted metabolomic analysis in the liver of mice exposed to A) 2 mg/week polystyrene B) 4 mg/week polystyrene, C) 2 mg/week mixed plastics, or D) 4 mg/week mixed plastics. Data plotted as log(2) fold change (p < 0.05). E) Venn diagram representing the significantly changed metabolites following microplastic exposures (p < 0.05 as compared to control). Exposures (5 µm microspheres): 2 x week for 4 weeks oral gavage; Mixed plastics: polystyrene, polyethylene, and poly-(lactic-co-glycolic acid); n=8 per group.

To identify the key metabolomic pathways altered in the liver following oral MP exposure, metabolomic pathway analysis was again performed. When comparing the impact on liver metabolomic pathways between the 2 mg and 4 mg/week polystyrene exposed groups, we observed significant changes (p < 0.05) in both PS groups in the following pathways: i) alanine, aspartate, and glutamate metabolism, ii) beta-alanine metabolism, iii) D-Glutamine and D-glutamate metabolism, iv) nitrogen metabolism, v) purine metabolism, and vi) tryptophan metabolism (Figure 6A, B). The beta-alanine and purine metabolic pathways were also increased in the 2mg/week and 4mg/week mixed plastic groups, respectively (Figure 6C, D). The 2mg/week and 4mg/week mixed plastic groups showed overlap in the i) amino acetyl-tRNA biosynthesis pathway, ii) propanoate metabolism, and iii) sphingolipid metabolism pathway (Figure 6C, D). The 2mg/week and 4mg/week mixed plastic groups also showed an increase in steroid biosynthesis and steroid metabolism, respectively. Interestingly, amongst all MP treated groups there were several pathways associated with amino acids biosynthesis or metabolism that was differentially regulated when compared to untreated mice.

**Figure 6.**
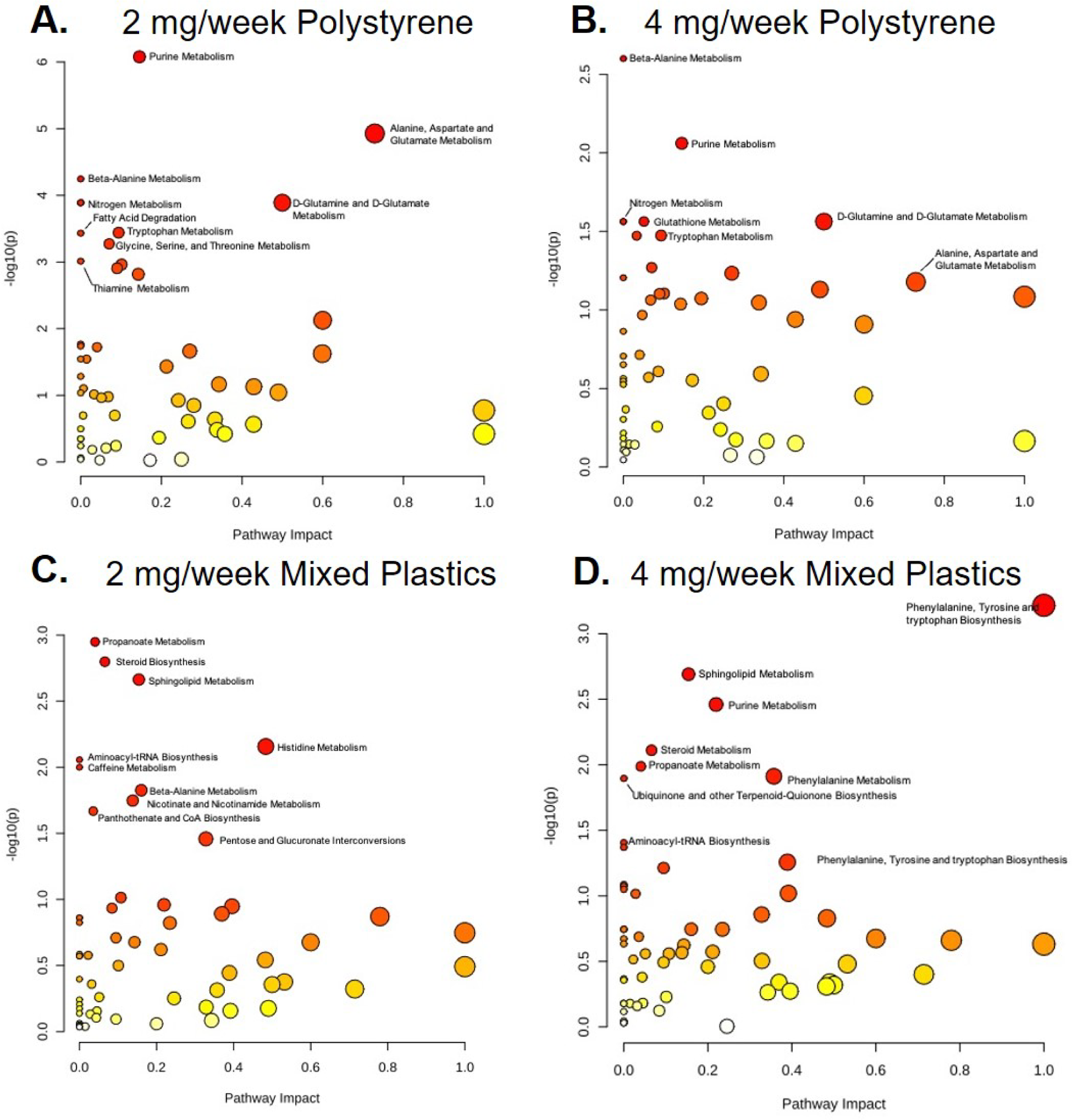
Metabolomic Pathway Analysis of alterations in the liver following oral MP exposure in mice exposed to A) 2 mg/week polystyrene, B) 4 mg/week polystyrene, C) 2 mg/week Mixed Plastics, or D) 4 mg/week Mixed Plastics. Exposures (5 µm microspheres): 2 x week for 4 weeks oral gavage; Mixed plastics: polystyrene, polyethylene, and poly-(lactic-co-glycolic acid); p < 0.05 as compared to control; n=8 per group.

Livers were next examined for metabolites associated with the nicotinate and nicotinamide metabolic pathway via targeted metabolomics. Liver samples from animals exposed to only polystyrene microspheres showed a shared upregulation of Nicotinic Acid Mononucelotide (NMN) and downregulation of 5-Hydroxy indoleacetic acid (HIAA) and L-Kynurenine in the 2 mg/week exposure and 4 mg/week exposure groups (Suppl. Figure 3A, B). The mixed plastic-exposed animals showed metabolite changes in both exposure groups, however, the metabolites being significantly upregulated between these groups were not comparable. In the 2 mg/week mixed plastic exposure group, melatonin was upregulated and 5-hydroxytryptophan and HIAA were downregulated (Suppl. Figure 3C). In contrast, the 4 mg/week mixed plastics exposure group showed a significant upregulation in metabolites nicotinamide riboside, 3-hydroxykynureneine, nicotinic acid adenine dinucleotide, and NAD and a significant downregulation of N’-formylkynurenine, tryptophan, and indole-3-lactic-acid (Suppl. Figure 3D). Taken together, both untargeted and targeted metabolomics show oral-gastric exposure of mixed and single MP leads to alterations in various metabolic pathways in the liver.

### Oral gastric exposure of PS-MP and mixed plastics alter the brain metabolome

Several studies have begun to show exposure to a single-type of MP or NP after ingestion can have far-reaching effects including alterations in the brain (Jin et al. 2022; Liang et al. 2022; Yang et al. 2023; Yang et al. 2022; Yin et al. 2022). To compare PS versus mixed plastic exposure, the prefrontal cortex of the brain was collected to perform untargeted metabolomics. Volcano plots for each exposure group showed patterns of metabolite changes within each group (**Figure 7A-D**). We observed that 33 metabolites in the polystyrene exposed group and 50 metabolites in the mixed plastic exposed group were significantly changed due to MP exposure (p < 0.05). We also observed that 12 (36.4%) metabolites were uniquely altered in the 2 mg/week polystyrene group and 18 (54.5%) metabolites were uniquely altered in the 4 mg/week polystyrene group, with 3 (9.1%) metabolites significantly altered in both dose groups (**Figure 7E**). In contrast, the mixed plastics exposure groups showed a much higher metabolomic alteration response, with 3 (6%) uniquely altered metabolites in the 2 mg/week group and 37 (74%) uniquely altered metabolites in the 4 mg/week group, and 10 (20%) metabolites shared between both of the mixed plastic groups.

**Figure 7:**
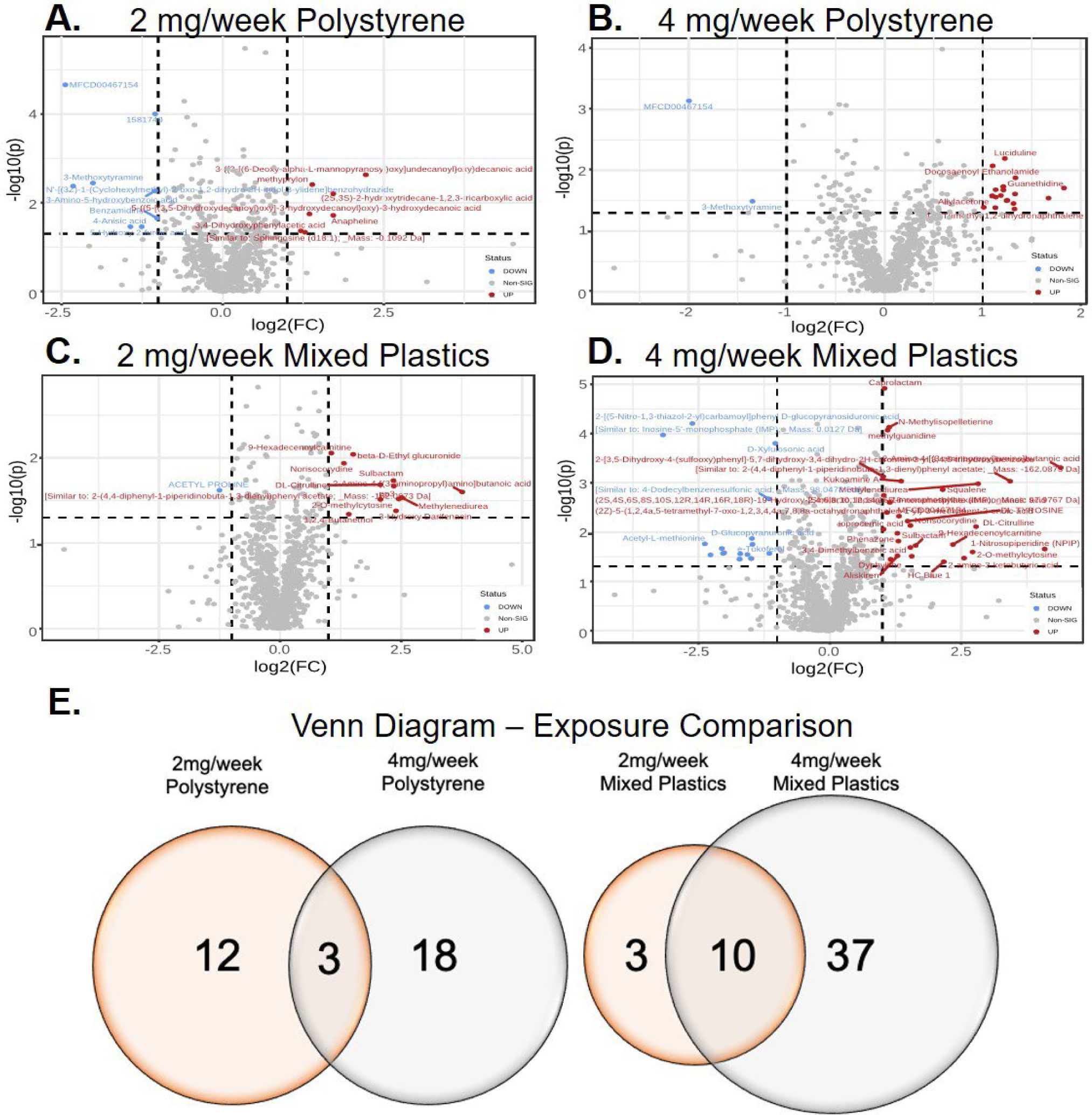
Untargeted metabolomic analysis of brain isolates from mice exposed to A) 2 mg/week polystyrene B) 4 mg/week polystyrene, C) 2 mg/week mixed plastics, or D) 4 mg/week mixed plastics. Data plotted as log(2) fold change (p < 0.05). E) Venn diagram representing the significantly changed metabolites following microplastic exposures (p < 0.05 as compared to control). Exposures (5 µm microspheres): 2 x week for 4 weeks oral gavage; Mixed plastics: polystyrene, polyethylene, and poly-(lactic-co-glycolic acid); n=8 per group.

To investigate the key metabolomic pathways altered by MP effects on the brain, metabolomic pathway analysis was again performed. When comparing the brain isolates from 2 mg and 4 mg/week polystyrene only exposed samples, we observed significant modulation (p < 0.05) of the following pathways: i) cysteine and methionine metabolism, ii) glycine, serine, and threonine metabolism, iii) sphingolipid metabolism, iv) tyrosine metabolism, and v) the xenobiotic metabolism regulated by cytochrome P450 (Figure 8A, B). Only the xenobiotic metabolism by cytochrome P450 pathway was shared by the 4 mg/week mixed plastic exposure group. Whereas both mixed plastic exposure groups shared a significant modulation in i) glycerolipid metabolism and ii) the steroid biosynthesis pathway (Figure 8C, D). Both 2mg/week exposure to PS or mixed plastics displayed an upregulation in metabolites associated with the i) D-glutamine and D-glutamate metabolism and ii) nitrogen metabolism. The 4mg/week exposure to PS or mixed plastics showed a significant increase in metabolites associated with the valine, leucine, and isoleucine degradation pathway.

**Figure 8.**
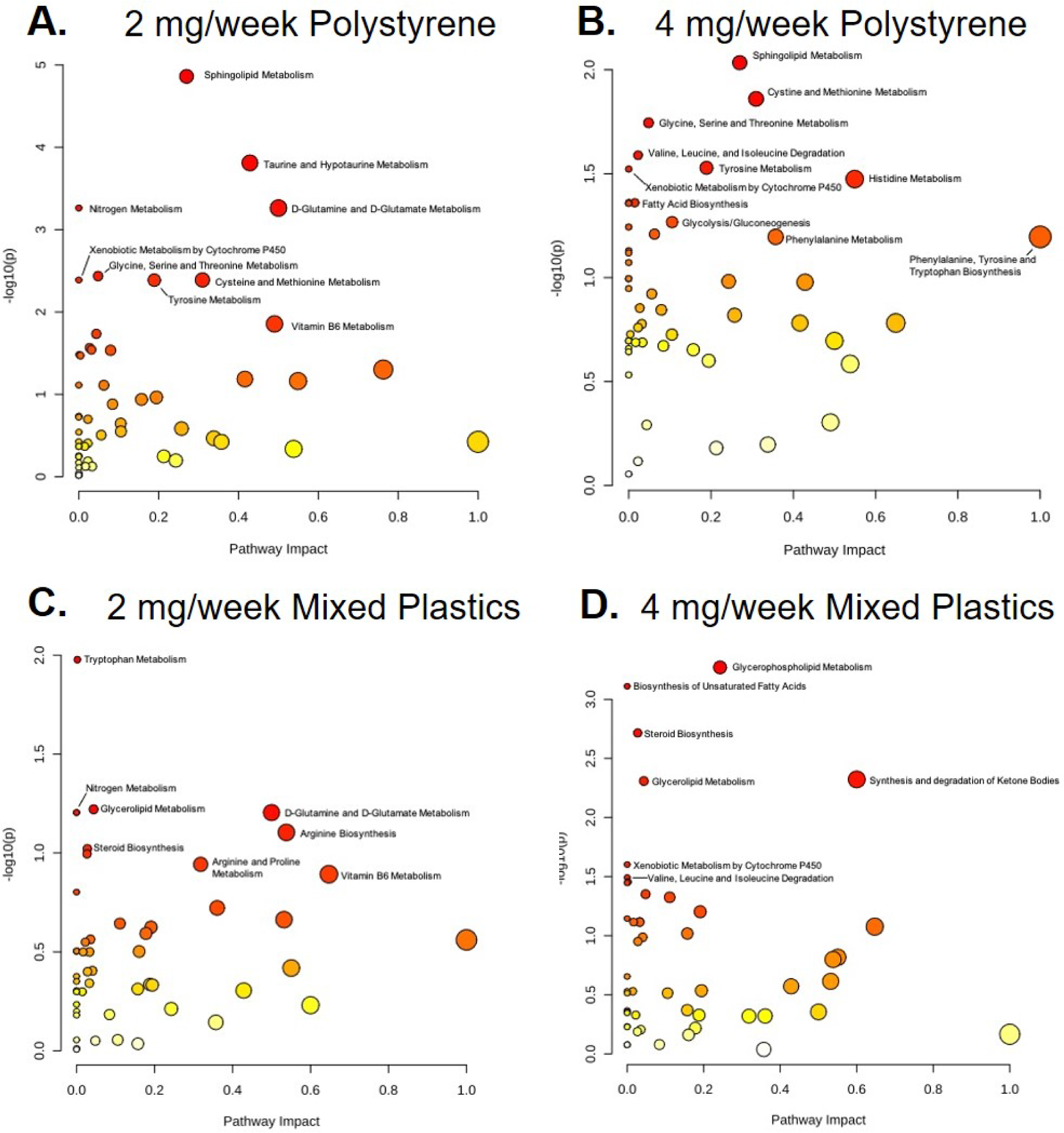
Metabolomic Pathway Analysis of alterations in the brain following oral MP exposure in mice exposed to A) 2 mg/week polystyrene, B) 4 mg/week polystyrene, C) 2 mg/week mixed plastics, or D) 4 mg/week mixed plastics. Exposures (5 µm microspheres): 2 x week for 4 weeks oral gavage; Mixed plastics: polystyrene, polyethylene, and poly-(lactic-co-glycolic acid); p < 0.05 as compared to control; n=8 per group.

When performing targeted metabolomic analysis in the brain for metabolites associated with the nicotinate and nicotinamide metabolic pathway we found animals exposed to polystyrene only showed upregulation of quinolinic acid and downregulation of nicotinic acid and tryptamine in the 2 mg/week exposure while the 4 mg/week dose group did not show any significant changes in upregulation or downregulation of metabolites (**Suppl. Figure 4A, B**). However, we did see trends of metabolites such as melatonin and nicotinic acid being downregulated and quinolinic acid and ADP ribose being upregulated, but these were not always significant. The mixed plastics exposed group showed multiple metabolite changes in both dose groups, with NADH being upregulated and L-kynurenine, HIAA and 3-hydroxykynurenine being significantly downregulated in both the 2 mg/week and 4 mg/week exposure groups (**Suppl. Figure 4C, D**). Overall, comparing mixed plastics versus PS alone demonstrates MP ingestion can have profound systemic effects in mammals including the brain.

## DISCUSSION

There is no doubt that all living organisms are being exposed to microplastics. While the physiological effects of MP pollution on marine organisms are well-documented (Auta et al. 2017; Barnes et al. 2009; Huerta Lwanga et al. 2017; Setala et al. 2014), the impacts on terrestrial organisms including humans are only beginning to be elucidated. The most common route of exposure appears to be through our diet (K. D. Cox et al. 2019; Huerta Lwanga et al. 2017; Kosuth et al. 2018; Liebezeit and Liebezeit 2014; Setala et al. 2014). Inhalation intake can also contribute to gut exposure through contaminated mucus ingestion (Beamish et al. 2011; Fournier et al. 2020; Lomer et al. 2002; Möller et al. 2004). To support this, there have been numerous reports showing in mammals and other species that MPs can accumulate in the gastrointestinal (GI) tract. Moreover, recent studies from various organisms including human blood and lungs suggest that MP accumulate can pass the GI tract (Jenner et al. 2022; Leslie et al. 2022). Furthermore, this systemic MP accumulation could drastically be increased in individuals with underlying conditions especially those that show signs of increased intestinal permeability such as inflammatory bowel disease (IBD), celiac disease, obesity, and non-alcoholic fatty liver disease (NAFLD) (Buning et al. 2012; Munkholm et al. 1994; Portincasa et al. 2021; Teshima et al. 2012; Turpin et al. 2020; Visser et al. 2009).

Until now, studies have only exposed mice to a single-type of MP and then performed metabolomics on either the serum, liver, or stool (Deng et al. 2017; Jin et al. 2019; Schwabl et al. 2019; Shi et al. 2022; Wang et al. 2022; Wright and Kelly 2017). However, humans are being exposed to a plethora of plastics and the assessment of mixed plastic exposure in animal models is critical to understand the true effects of plastic pollution and health outcomes. Our findings provide further support that MP can become embedded in other internal organs after ingestion. After oral gastric MP exposure in a healthy mouse, we find MPs can pass through the digestive system and translocate via the systemic circulation to distant organs (i.e., brain, liver, and kidney). Moreover, we show that a four-week PS alone or mixed plastic exposure can impact various metabolomic pathways in the colon, liver, and brain of mice when compared to unexposed mice. The most common pathways dysregulated in the colon, liver and brain were associated with amino acids. Many other metabolites such as purines and pyrimidines and neurotransmitters such as glutamate are products of amino acid metabolism that we see alterations in metabolic pathways associated with purines, pyrimidines, and glutamate in our MP-exposed mice. Amino acids are fundamental for human health as they influence numerous physiological processes and disruptions in amino acid metabolism have been linked to numerous inflammatory and metabolic diseases (Gaggini et al. 2018; Grohmann et al. 2017; Handzlik et al. 2023; Hasegawa et al. 2020; Lamichhane et al. 2019).

Interestingly, our metabolomics data not only showed differences in the colon, liver and brain metabolome when comparing our mixed plastic exposure to PS alone groups, but also showed a difference when examining the two doses for each group. This became more apparent when we performed targeted metabolomics on the metabolites associated with the nicotinate and nicotinamide metabolic pathway. The greatest impact on the host metabolome was in the following order - colon, liver, and then brain. One reason for these fewer metabolomic changes in the brain as compared to the colon and liver may be due to the overall greater accumulation of MPs in these tissues and the crosstalk between the gut and liver. Other reasons for more extensive metabolomic alterations occurring in the colon and liver may be due to the role that these tissues play in overall break down, digestion, detoxification and synthesis of consumed products.

In this study, the prefrontal cortex was the only region of the brain evaluated. Thus, we cannot rule out that other portions of the brain may accumulate MP and this could cause further changes in the brain metabolome. Nevertheless, one alteration of interest was the modulation of the xenobiotic pathway by cytochrome P450 seen in the brain of animals exposed to 4mg/week of PS and mixed plastics. Although metabolism is primarily carried out in the liver, xenobiotic metabolism by cytochrome P450 alteration in the brain as a result of MP exposure could point to potential neurotoxic effects. This data supports the various studies showing MPs can be neurotoxic (Jin et al. 2022; Liang et al. 2022; Yang et al. 2023; Yang et al. 2022; Yin et al. 2022). Furthermore, it suggests that ingestion of MPs over time could cause adverse neurodevelopmental outcomes or trigger the development of neurodegenerative diseases (Silva-Adaya et al. 2021). While this finding requires further investigation, the effects of MPs on our central nervous system may be an interesting avenue for future research.

We hope this study will be used as a model to explore chronic exposure outcomes associated with mixed plastic exposure. Thus, leading to novel techniques to identify their potential risks on human health and future quantitative method development to establish MP-associated metabolite presence. Future studies will focus on novel techniques to adequately identify and quantify specific plasticizers and to investigate the functional impacts of altered metabolites due to systemic uptake and distribution of MPs. Taken together, our data highlight potential risks that the different types, mixtures, and doses of MP exposure can impact health outcomes.

### Limitation of study

Currently, there is substantial research on possible connections of MPs exposure and poor health outcomes in wildlife, but there has been limited investigation into long-term human health outcomes (Vethaak and Legler 2021). When ingested, MPs have the potential to expose organisms to higher concentrations of monomers, polymers, or chemicals associated with the manufacturing process that could potentiate their toxicity (Prata et al. 2020). It is believed the plastics living organisms are ingesting contain chemicals that can further exacerbate plastic-associated toxicity that has been detailed (Chang et al. 2021; Yong et al. 2020). The plastics utilized in this study are commercially bought and do not contain chemicals such as phthalates, Bisphenol A (BPA), polyfluorinated alkyl substances (PFAs) (Arrigo et al. 2023; Barhoumi et al. 2022; Sharma et al. 2022). These chemical additives could an extra layer of problems but we believe our study shows that MP have far-reaching effects after ingestion. An although there is still ongoing research to identify and understand the widespread human health risk of MPs, the current study helped to identify potential organ-specific metabolomic pathway alterations that are associated with different types, mixtures, and doses of MP. Further investigation will need to be performed to identify if these metabolomic alterations may play a role in inflammation, immune regulation, metabolism, multiorgan dysfunction, and even potentially exacerbate conditions such as IBD, NAFLD, and obesity (Horn et al. 2021).

### AUTHOR CONTRIBUTION

MMG and ASR performed all analysis, tissue collection, and isolation with the help from SDM, JLMH, CF, EEH, DPS, RT, JGE, AB, RPH, SL, GH, KJK, JYC, RG, and JGI. HG and YJ contributed to metabolomics and analysis. Exposures were performed by MMG, ASR, SL, and GH. MMG, MJC, and EFC participated in writing the manuscript. MMG, JGI, MJC and EFC designed the study, analyzed data and wrote the paper. All authors approved the final version of the manuscript.

### GRANT FUNDING

Funding was supported in part by the National Center for Research Resources and the National Center for Advancing Translational Sciences of the National Institutes of Health (NIH) through grant no. UL1TR001449 (EFC) in part by NIH grant 1R01 ES032037-01A1 (EFC), in part by NIH grant P20GM121176 (EFC), and P20GM130422 (MJC). and in part by NIH grant K12GM088021 (M.G.; ASERT-IRACDA), in part by NIMHD grant P50MD015706 (JGI), in part by pilot funding from P20GM130422 (EEH), and in part by P30CA118100.

### COMPLIANCE WITH ETHICAL STANDARDS

Studies were conducted with full approval by the Institutional Animal Care and Use Committees of the University of New Mexico.

## ACKNOWLEDGMENTS

We would like to acknowledge Jesse Benson Hesch for her valuable contribution to the editing of this manuscript. Her expertise and meticulous attention to detail significantly enhanced the quality of our work, and preparation of this manuscript.

---

**Supplemental Figure 1.**
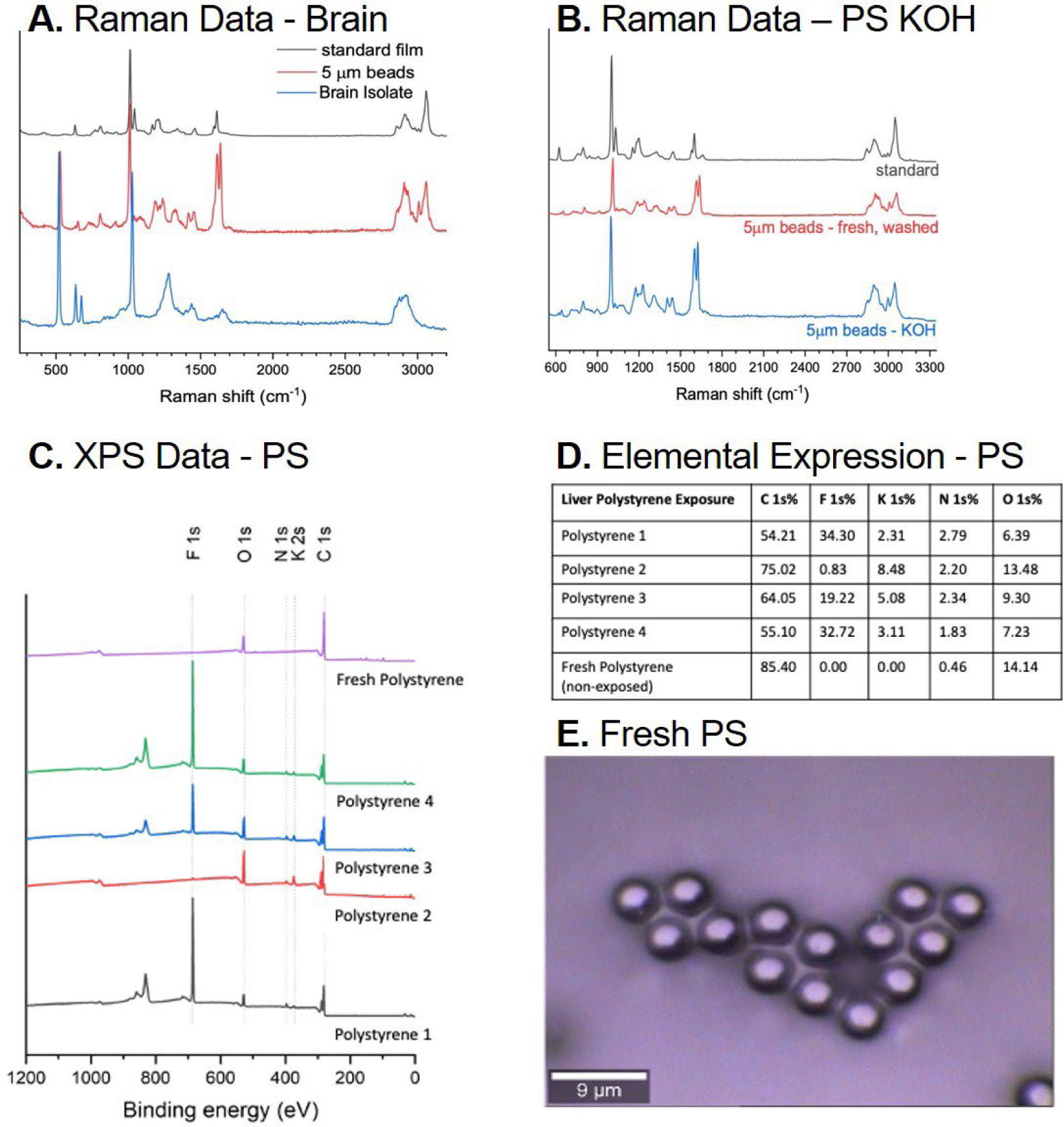
Raman Spectroscopy confirms identification of polystyrene microspheres in the brain and XPS analysis reveals presence of microplastics in the liver. A) Raman spectra of polystyrene microspheres isolated from brain isolates compared to fresh 5 µm polystyrene microspheres and standard library film polystyrene. Brain tissue was isolated from mice exposed twice a week for 4 weeks to 2 or 4 mg 5 µm polystyrene microspheres. B) Raman Spectroscopy of 5 µm polystyrene microspheres exposed to potassium hydroxide (KOH) digestion compared to fresh 5 µm polystyrene microspheres and standard library film polystyrene. C) XPS data comparing 4 representative polystyrene microspheres isolated from the liver of mice exposed twice a week for 4 weeks via oral gavage to fresh polystyrene microspheres at both 2mg/week and 4mg/week exposure groups. Polystyrene 1 and Polystyrene 2 were representative of mice exposed to polystyrene at the 2mg/week dose. Polystyrene 3 and Polystyrene 4 were representative of mice exposed to polystyrene at the 4mg/week. D) Elemental expression of each polystyrene sample isolated from the liver of exposed mice compared to fresh polystyrene microspheres. E) Visualization of fresh polystyrene microspheres used for XPS Analysis (60x magnification).

**Supplemental Figure 2.**
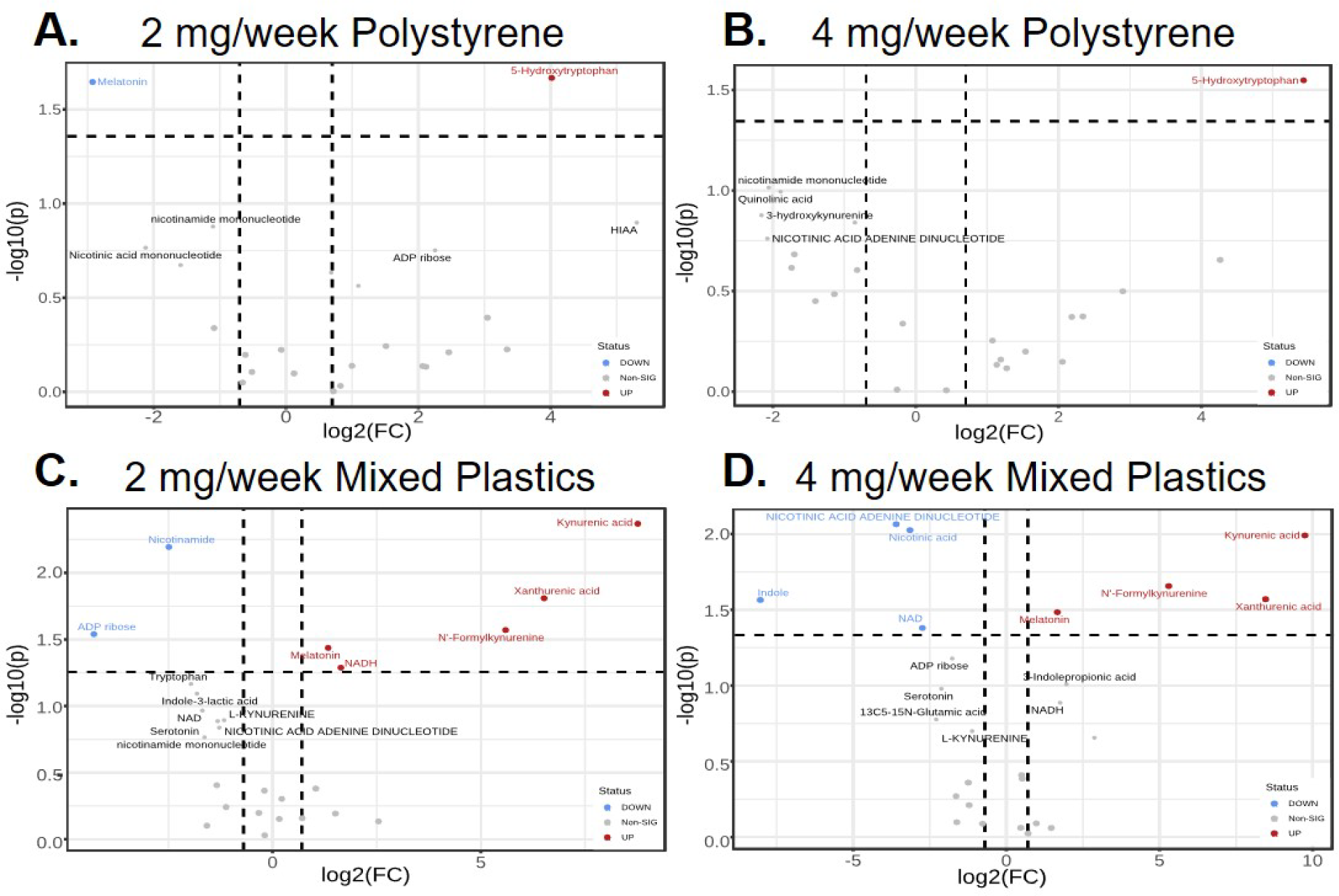
Volcano plots of targeted metabolomic analysis in the colon of mice exposed to A) 2 mg/week polystyrene (5 µm), B) 4 mg/week polystyrene, C) 2 mg/week mixed plastics (5 µm), and D) 4 mg/week mixed plastics. Exposures were four weeks in duration twice weekly via oral gavage. Mixed plastics: polystyrene, polyethylene, and poly-(lactic-co-glycolic acid) microspheres. Exposure groups were compared to 0 mg/week controls. Data are plotted as log(2) fold change (p < 0.05; n=8 per group).

**Supplemental Figure 3.**
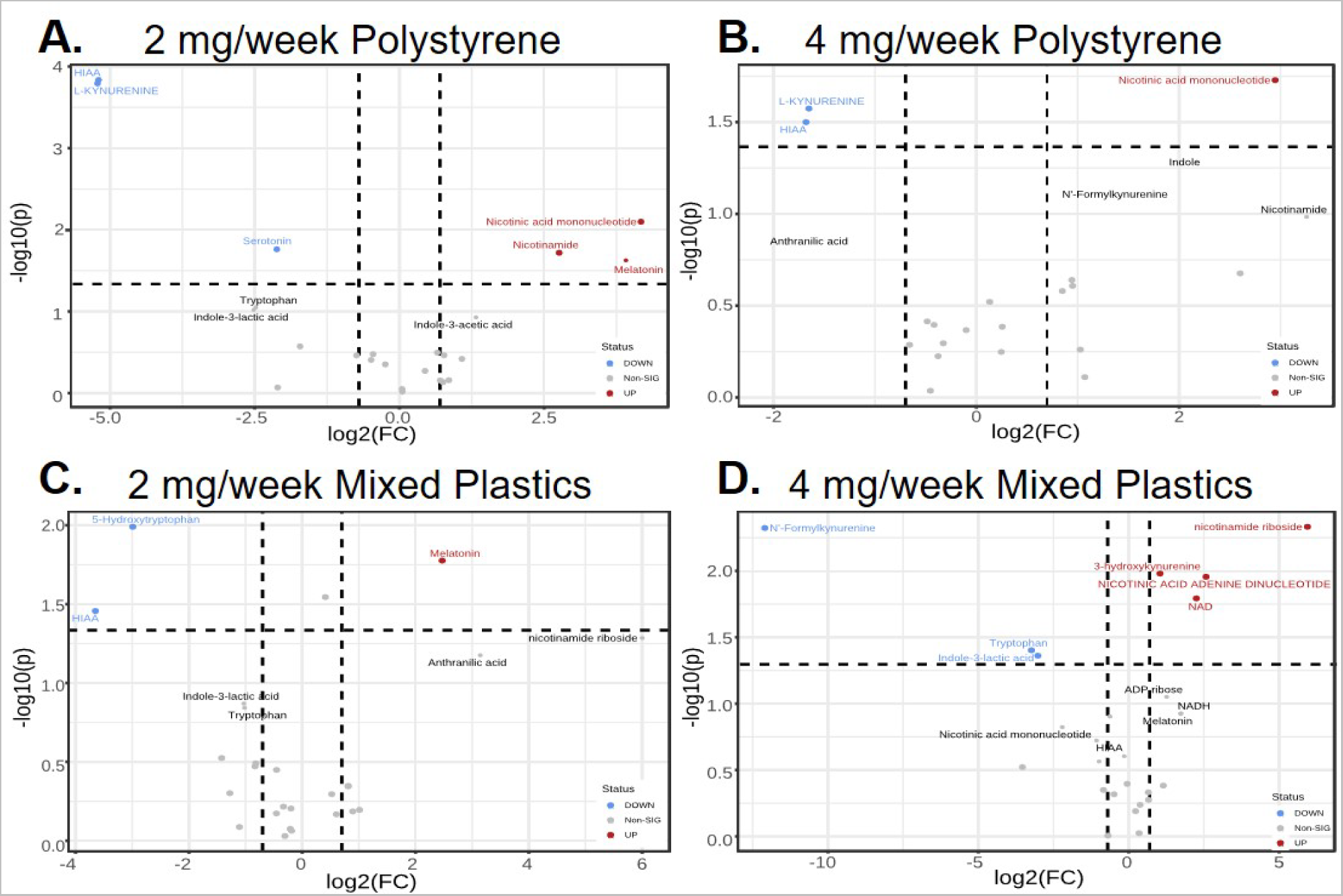
Volcano plots showing targeted metabolomic analysis of livers from mice exposed to A) 2 mg/week polystyrene, B) 4 mg/week polystyrene C) 2 mg/week mixed plastics, or D) 4 mg/week mixed plastics. Exposures were four weeks in duration twice weekly via oral gavage. Mixed plastics: polystyrene, polyethylene, and poly-(lactic-co-glycolic acid) microspheres. Exposure groups were compared to 0 mg/week controls. Data are plotted as log(2) fold change (p < 0.05; n=8 per group).

**Supplemental Figure 4.**
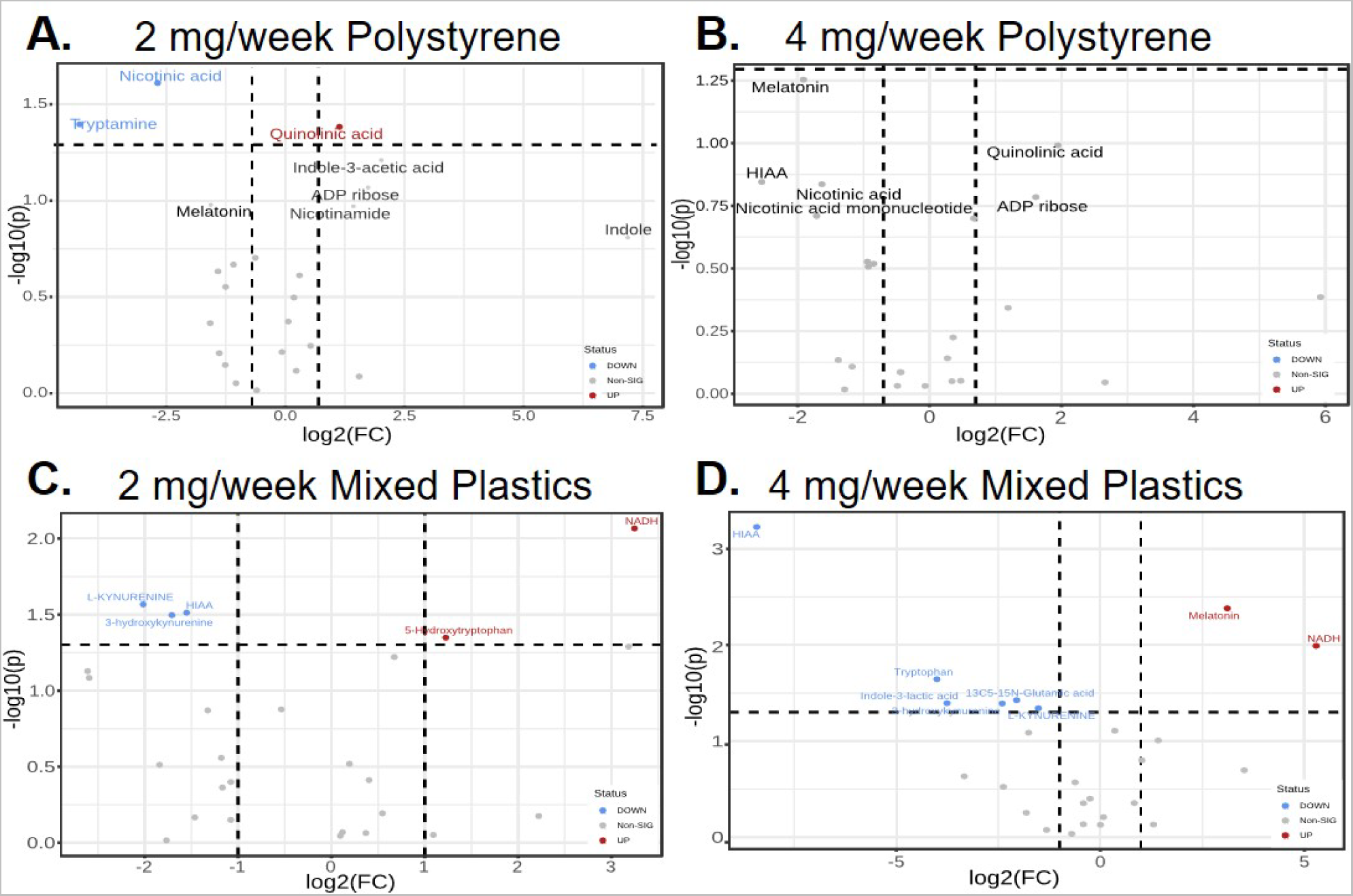
Volcano plots showing targeted metabolomic analysis of brain isolates from mice exposed to A) 2 mg/week polystyrene, B) 4 mg/week polystyrene C) 2 mg/week mixed plastics, or D) 4 mg/week mixed plastics. Exposures were four weeks in duration twice weekly via oral gavage. Mixed plastics consisted of polystyrene, polyethylene, and poly-(lactic-co-glycolic acid) microspheres. Exposure groups were compared to 0 mg/week controls. Data are plotted as log(2) fold change (p < 0.05; n=8 per group).

